# *In situ* relationships between microbiota and potential pathobiota in *Arabidopsis thaliana*

**DOI:** 10.1101/261602

**Authors:** Claudia Bartoli, Léa Frachon, Matthieu Barret, Mylène Rigal, Carine Huard-Chauveau, Baptiste Mayjonade, Catherine Zanchetta, Olivier Bouchez, Dominique Roby, Sébastien Carrère, Fabrice Roux

## Abstract

A current challenge in microbial pathogenesis is to identify biological control agents that may prevent and/or limit host invasion by microbial pathogens. *In natura,* hosts are often infected by multiple pathogens. However, most of the current studies have been performed under laboratory controlled conditions and by taking into account the interaction between a single commensal species and a single pathogenic species. The next step is therefore to explore the relationships between host-microbial communities (microbiota) and microbial members with potential pathogenic behavior (pathobiota) in a realistic ecological context. In the present study, we investigated such relationships within root and leaf associated bacterial communities of 163 ecologically contrasted *Arabidopsis thaliana* populations sampled across two seasons in South-West of France. In agreement with the theory of the invasion paradox, we observed a significant humped-back relationship between microbiota and pathobiota α-diversity that was robust between both seasons and plant organs. In most populations, we also observed a strong dynamics of microbiota composition between seasons. Accordingly, the potential pathobiota composition was explained by combinations of season-specific microbiota OTUs. This result suggests that the potential biomarkers controlling pathogen’s invasion are highly dynamic.

## INTRODUCTION

In the last decade, a conspicuous effort has been made to characterize and understand the role of host-associated microorganisms. Animals and plants are currently considered as holobionts associated with the commensals that inhabit and evolve with them (Youle *et al.*, 2013; Bordenstein and Theis, 2015; Vandenkoornhuyse *et al.*, 2015). However, how commensal microbes interact with pathogens remains largely unknown. It has been recently reported that the human gut microbiota can protect against the potential overgrowth of indigenous opportunistic pathobionts (i.e. pathogenic species that inhabit the host at a low bacterial population size) (Vayssier-Taussat *et al.*, 2014) or pathogenic invaders (Kamada *et al.*, 2013) by niche competition and/or induction of the host immune system (Belkaid and Hand, 2014). In plants, both rhizosphere-and phyllosphere-associated bacteria can directly or indirectly increase host resistance to phytopathogenic microorganisms *via* production of antimicrobial compounds or elicitation of plant defences (Mendes *et al.*, 2013; Berg *et al.*, 2014; Bodenhausen *et al.*, 2014; Haney *et al.,* 2015; Santhanam et *al.,* 2015; Ritpitakphong *et al.*, 2016). For example, plant-associated strains belonging to the *Sphingomonas* genus can protect the model annual plant *Arabidopsis thaliana* from infections caused by the causal agent of bacterial spot *Pseudomonas syringae* (Innerebner *et al.*, 2011). Although these studies support the importance of certain microbes in protecting plants against infections, they mainly focused on the interaction between a single commensal species and a single pathogenic strain. Nevertheless, as previously observed in animals (Singer, 2010; Short *et al.*, 2014), plants are often infected *in natura* by multiple pathogens (Bartoli *et al.*, 2016; Lamichhane and Venturi, 2015; Roux and Bergelson, 2016). Therefore, the current challenge in plant pathology is to investigate the relationships between resident microbial communities and pathogenic microbes at the community level, i.e. between microbiota (defined here as the microbial communities inhabiting plants) and pathobiota (defined here as the complex of microorganisms with the potential to cause disease on a given plant host; Bartoli and Roux, 2017).

Understanding how the native cortege of beneficial/commensal microbes interacts to protect plants against the invasion of a cortege of pathogenic species requires testing for community ecological theories (Mallon, *et al.*, 2018). In plant community ecology, substantial efforts have been made in the last decades to understand the relationships between α-diversity of resident species and α-diversity of invasive species (Fridley et al. 2007) by both applying spatial pattern studies and constructed community studies (Naeem et al. 2000; Tomasetto et al. 2013). In a large number of studies conducted at large spatial scale, species richness of both resident and invasive species were found to be positively correlated with the quality of the abiotic environment, thereby leading to a positive relationship between resident species diversity and invader success (Eisenhauer *et al.*, 2013a) (Supplementary Figure S1). On the other hand, as predicted by Elton’s theory (1958), constructed community studies performed in homogenous abiotic environments revealed a negative relationship between resident species diversity and invader success (Naeem et al., 2000). This negative relationship is explained by the negative impact of the increased resident species diversity on resource availability, which is in turn detrimental for the establishment of the invasive species (Supplementary Figure S1). These two contrasted phenomena lead to the invasion paradox (humped-back relationship between the diversity of invaders and the diversity of resident species, Supplementary Figure S1), which is therefore predicted to mainly result from the interplay between species diversity, resource availability and niche dimensionality in ecologically relevant conditions (Mallon *et al.,* 2015a).

Testing for the invasion paradox between microbiota and pathobiota therefore requires a characterization of the microbial communities across the range of native habitats encountered by a given plant species (Roux & Bergelson 2016). Accordingly, recent studies highlighted that the ecology of the habitats where both plants and microorganisms co-exist and evolve is a crucial variable to take into account when investigating the relationships between a host and its microbes participating in the holobiont system (Agler *et al.* 2016, Wagner *et al*. 2016). In addition, in native habitats, plants are naturally exposed to pathogens whose attacks are influenced by local abiotic/biotic conditions (Bartoli *et al.*, 2016).

In this study, we aimed to investigate the *in situ* relationships between bacterial communities and the potential bacterial pathobiota in 163 natural *A. thaliana* populations collected in southwest of France and inhabiting ecologically contrasted habitats. In particular, we aimed (i) to test for the invasion paradox and (ii) to identify the combinations of microbial species that can prevent and/or limit pathogen invasion. Because bacterial communities associated with plants can rapidly change within the host life cycle (Chaparro *et al.*, 2014; Schlaeppi *et al.*, 2014; Copeland *et al.*, 2015; Agler *et al.*, 2016) and can largely differ between plant compartments (Bodenhausen *et al.*, 2013; Coleman-Derr et al., 2016; Wagner et *al.,* 2016), we described both microbiota and potential pathobiota in the leaf and root compartments across two seasons within a single life cycle of *A. thaliana*.

## MATERIAL AND METHODS

### Identification of *A. thaliana* populations

In this study, we focused on 163 natural *A. thaliana* populations identified in May 2014 in the Midi-Pyrénées region (Figure 1a, Supplementary Table 1). These populations were chosen to maximize the diversity of habitats (such as climate, soil type, vegetation type and degree of anthropogenic perturbation) encountered by *A. thaliana* (Figure 1b). Because the 163 populations strongly differed in their main germination cohort in autumn 2014 (early November *vs* early December; Supplementary Text), we defined three seasonal groups hereafter named (i) ‘autumn’ corresponding to 84 populations collected in November/December 2014, (ii) ‘spring with autumn’ corresponding to 80 populations already sampled in autumn and additionally sampled in early-spring (February/March 2015), and (iii) ‘spring without autumn’, corresponding to 79 populations only sampled in early-spring (February/March 2015) (Supplementary Text).

**Figure 1.**
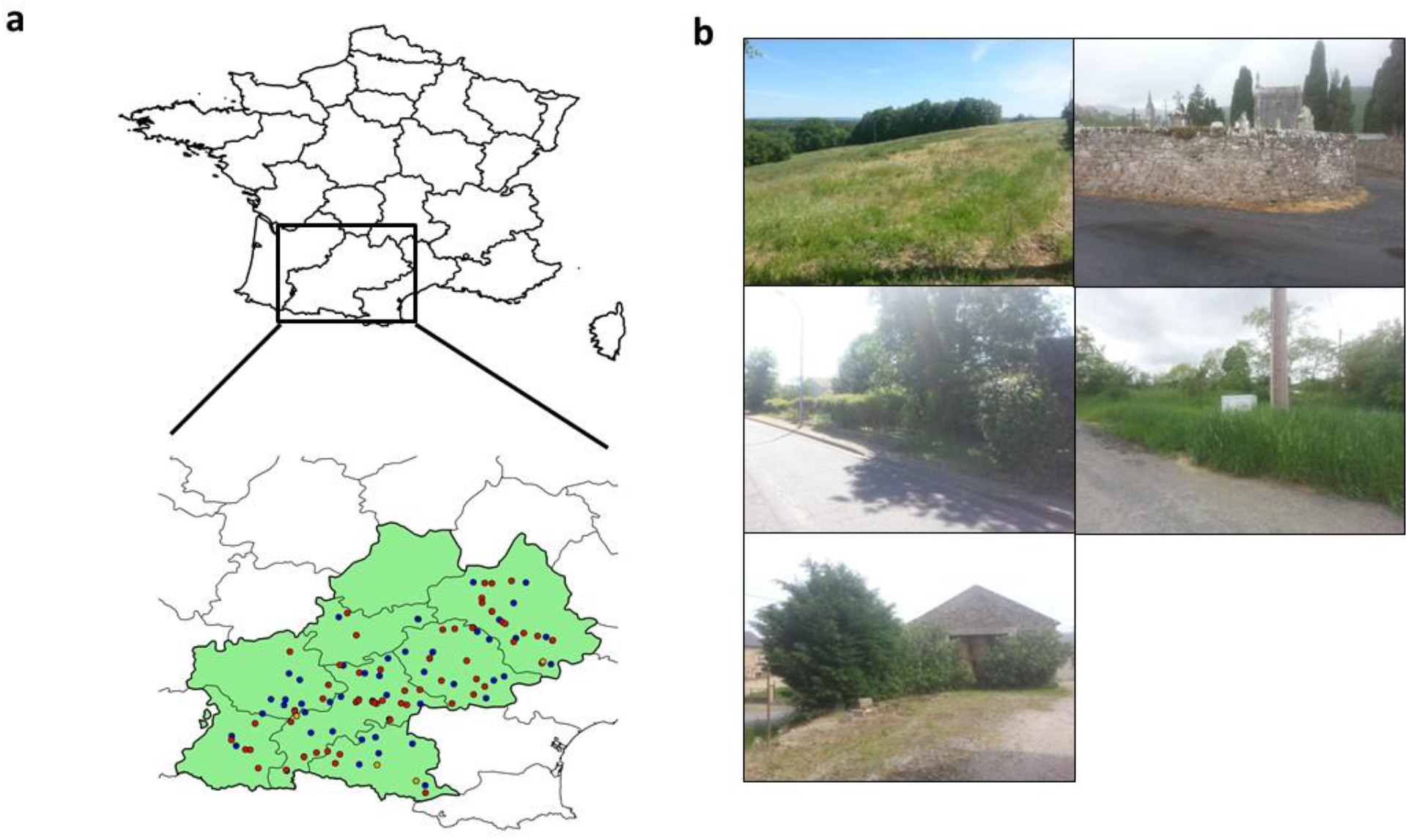
Plant material. (**a**) Location of the 163 *A. thaliana* populations across the Midi-Pyrénées region (southwest of France). The average distance among populations was 99.9 km (median = 92.6 km, SD = 55.4 km). Blue dots represent the 80 populations collected in both autumn and spring, red dots represent the 79 populations collected in spring only and orange dots represent the 4 populations collected in autumn only. The map clearly shows that the range of sampling across the Midi-Pyrénées region was homogeneous during the two sampling seasons. (**b**) Diversity of habitats encountered by *A. thaliana* in the Midi-Pyrénées region.

To avoid a confounding effect between the sampling date and geographical origin, populations were randomly collected during the sampling periods in autumn 2014 and early-spring 2015.

### Sampling, generation of the *gyrB* amplicons and sequences

The bacterial communities were characterized through amplification of a fraction of *gyrB* gene encoding for the bacterial gyrase *β* subunit. This molecular marker has a deeper taxonomic resolution than other molecular markers designed on the hypervariable regions of the 16S rRNA gene (Barret *et al.*, 2015), thereby allowing to distinguish bacterial OTUs at the species level. Furthermore, this single-copy gene limits the overestimation of taxa carrying multiple copies of *rrn* operons. In this study, based on synthetic communities, we confirmed a better taxonomic resolution of the *gyrB* gene compared with the 16S rRNA gene (Supplementary Text, Supplementary Data 1).

To characterize the bacterial communities, ∼4 individuals per population and per season were sampled *in situ* at the rosette stage resulting in a total number of 1,912 leaf and root samples (Supplementary Text). The epiphytic and endophytic bacterial components of either leaf or root samples were not separated. For each plant compartment, we therefore extracted the total DNA of both epiphytic and endophytic microbes (Supplementary Text). The amplicon *gyrB* was amplified with some modification of the protocol described in Barret *et al.* (2015). In particular, three internal tags were added at each 5′ and 3′ of the original primers to allow the multiplexing of three 96-well plates (Supplementary Text).

For each sample, PCR amplifications were repeated three times and technical replicates were pooled in a unique PCR plate. PCR products were purified by using Agencourt^®^AMPure^®^ magnetic beads following manufacturer’s instructions and purified amplicons were quantified with Nanodrop and appropriately diluted to obtain an equimolar concentration. Two μl of equimolar PCR purified products were used for a second PCR with the Illumina adaptors. The second PCR amplicons were then purified and quantified as described above to obtain a unique equimolar pool. The latter was quantified by RQ-PCR and then sequenced with Illumina MiSeq 2X250 v3 (Illumina Inc., San Diego, CA, USA) in the GeT-PlaGe Platform (Toulouse, France). MS-102-3003 MiSeq Reagent Kit v3 600 cycle was used for this purpose.

### Bioinformatics analysis and data curation

Reads were demultiplexed by considering the three internal tags. After demultiplexing, the average number of sequences per sample was of 46,135 ± 19,820. Prior further analysis, the negative controls – consisted by the sterilized water used to clean up the samples during field sampling, the sterilized DNA free water used to elute the DNA after phenol/chloroform extraction and the DNA free water used for PCR amplifications – were checked for presence/absence of amplicons by blasting them against the *gyrB* database composed by 30,627 sequences (Barret *et al*. 2015; Supplementary Text). Because negative controls showed no trace of entire *gyrB* sequences, they were removed before clustering. Samples showing low sequence quality were also removed before clustering, resulting in a total of 1,903 samples. Taxonomic affiliation of *gyrB* sequences was performed by using a Bayesian classifier (Wang *et al.*, 2007) implemented in the classify.seqs command of mothur (Schloss *et al.*, 2009) against an in-house *gyrB* database containing 30,627 representative sequences with an 80% bootstrap confidence score. Clustering of sequences into Operational Taxonomic Units (OTUs) was performed with Swarm (Mahé *et al.*, 2015) by using a clustering threshold (*d*) = 1. Only the OTUs that were composed by a minimum of 5 sequences across all samples were kept, resulting in a total of 278,336 OTUs. Then, we applied two steps of filtering. First, to control for sampling limitation within each sample, we estimated a Good’s coverage score for each sample (Schloss *et al.*, 2009). Based on the distribution of the Good’s coverage score, only samples with a score more than 0.5 were considered. Second, only OTUs showing a minimum relative abundance of 1% in at least one sample were selected by using a home-made perl program. The final data set corresponded to a matrix of 1,655 samples by 6,627 OTUs. The 6,627 OTUs obtained after the filtering of the data represented 2.4 % of the totality of the OTUs but 55.6 % of the totality of the reads (Supplementary Figure S2). In addition, the mean number of reads per OTU maintained after filtering was 51.3 fold higher than the one of the discarded OTUs (Supplementary Figure S2). This final data set was used to determine the matrix composed by the potential pathogenic species. More precisely, the potential pathobiota in both leaves and roots of the 163 *A. thaliana* populations was determined by using a list of the phytopathogenic bacteria established by the International Society of Plant Pathology Committee on the Taxonomy of Plant Pathogenic Bacteria (ISPP-CTPPB; Supplementary Data 2) (Bull *et al.*, 2010, 2014). This potential pathobiota list (Supplementary Data Set 2) was composed by 199 bacterial species that were filtered on the microbiota matrix. The potential pathobiota matrix resulted in 11 bacterial species that were further curated. More precisely,we only considered: i) the bacterial species already reported to colonize *A. thaliana* (*Pseudomonas syringae*, *Pseudomonas viridiflava*, *Pantoea agglomerans, Sphingomonas melonis* and *Xanthomonas campestris*) (Jakob *et al.*, 2002, Kniskern *et al.*, 2007), ii) the bacterial species pathogenic on Brassicaceae species (*Pseudomonas marginalis*, *Streptomyces scabei* and *Xanthomonas perforans*) (Charron & Sams, 2002, Lerat *et al.*, 2009), iii) the causal agent of lamb’s lettuce spot *Acidovorax valerianellae* reported to be widely distributed in France (Gardan *et al.*, 2003). The bacterial species *Diaphorobacter oryzae* and *Janthinobacterium agaricidamnosum* were removed from the potential pathobiota matrix because they have been reported previously only in non-plant habitats (Pham *et al.*, 2009) and mushrooms (Lincoln *et al.*, 1999), respectively. The resulted potential pathobiota matrix was composed by 1203 samples and 29 OTUs characterizing the nine bacterial species listed above (i.e. a given bacterial species can be included in more than one OTU). The 29 potential pathogenic OTUs were removed from the microbiota matrix. Therefore, the final microbiota matrix used for further analysis was composed by 6,598 OTUs.

### Analysis of the α and β-diversity and characterization of the potential pathobiota

Shannon diversity and observed species richness were estimated on the final OTU matrix by using the summary.single function of mothur (Schloss *et al.*, 2009). Indexes of α-diversity were also calculated by sample rarefaction of 300/600/900 iters. Results between non-rarefied and rarefied samples were similar and the complete non-rarefied data set was used for the calculation of microbiota diversity.

Due to sparsity of the OTU matrix, a Hellinger transformation (Legendre and Gallagher, 2001) was performed by using the *vegan* R package (Jari *et al.*, 2009) and the relative Hellinger distance was inferred with the *decostand* command in the *vegan* R package prior β-diversity analyses. Following Ramette (2007), the resulting Hellinger distance matrix was reduced by running a Principal Coordinates Analysis (PCoA) with the ape R package (Paradis *et al.*, 2004). Because PCoA performed on Hellinger distance matrix based on rarefied data and on Jaccard similarity coefficient matrix led to similar patterns of ordination (Supplementary Figure S3), the PCoA coordinates from the non-rarefied Hellinger distance matrix were retrieved and used for statistical analysis described below.

Non-metric multidimensional scaling (NMDS) was also run on the Hellinger distance matrix. However, the values of stress were 0.431 and 0.293 for 2D and 3D NMDS ordination space, respectively. These stress values suggest a lack of fit between the ranks on the NMDS ordination configuration and the ranks in the original distance matrix (Ramette, 2007).

### Statistical analysis

Natural variation for the eight descriptors of microbiota (i.e. species richness, α-diversity Shannon index, first and second PCoA axes) and potential pathobiota microbiota (i.e. species richness, α-diversity Shannon index, first and second PCoA axes) was explored using different mixed models (Supplementary Text). A correction for the number of tests was performed to control the FDR at a nominal level of 5%.

In order to study the relationship between microbiota α-diversity and potential pathobiota α-diversity, linear and non-linear regressions were fitted using the ‘lm’ and ‘nls’ functions implemented in the *R* environment, respectively (Supplementary Text). Using a paired sample *t*-test, model selection was performed by comparing the goodness of fit between linear and non-linear models across the ‘diversity estimate x plant compartment x seasonal group’ combinations. In order to confirm the significance of the humped-back curve observed between diversity estimates (species richness and Shannon α-diversity) of the microbiota and the potential pathobiota, the parameters of the non-linear model were compared to a null distribution of those parameters obtained by creating 100 random microbiota OTU matrices paired with 100 random pathobiota OTU matrices (Supplementary Text).

Although pathogen-focused network analysis has been already used for investigating the relationships between whole microbiota and microbial pathogens, this method appears only suitable for studying monospecific interactions between a single pathogenic species and the rest of the microbial community members (Poudel *et al.*, 2016). In order to include higher-order interactions in the study of the relationship between microbiota and potential pathobiota composition, a sparse Partial Least Square Regression (sPLSR) (Lê Cao *et al.*, 2008; Carrascal *et al.*, 2009) was therefore adopted to maximize the covariance between linear combinations of relative abundances of OTUs from the microbiota (matrix X) and linear combinations of relative abundances of species from the potential pathobiota (matrix Y) (Supplementary Text). Significance of the OTUs included in the linear combinations was estimated by a Jackknife resampling approach by leaving out 10% of the samples 1,000 times (Supplementary Text).

### Data availability

The raw FastQ reads were deposited in the Sequence Read Archive (SRA) of NCBI https://trace.ncbi.nlm.nih.gov/Traces/sra/?study=SRP096011 under the study number SRP096011. The filtered matrix for both microbiota and potential pathobiota are available in Supplementary Data 4 and Supplementary Data 5 respectively. Raw values for both α and β-diversity used for statistical analysis are available in Supplementary Data 6.

## RESULTS

### Characterization of the *A. thaliana* microbiota and potential pathobiota

We sampled 163 natural *A. thaliana* populations chosen to maximize the diversity of habitats encountered by *A. thaliana* in the Midi-Pyrénées region (Figure 1). Due to differences in germination timing in autumn, about half of the populations were sampled both in autumn and in spring, whereas the other half of populations were sampled only in spring, thereby leading to three seasonal groups of populations, that is ‘autumn’, ‘spring with autumn’ and ‘spring without autumn’.

We obtained 18,610,383 high-quality reads across 1,655 samples, with on average ∼10,136 reads per sample. After data filtering, we identified 6,627 non-singleton bacterial OTUs. A large amount of these OTUs were specific to roots or leaves, as only ∼8.1% OTUs (n = 540 OTUs) were shared between both plant compartments. However, the relative abundance of OTUs shared between leaf and root samples was 20.2 and 16.0 higher than the relative abundance of leaf and root specific OTUs. This suggests that generalists OTUs are dominant members of the *Arabidopsis thaliana* microbiota.

As commonly observed in *A. thaliana* and other plant species (Lundberg *et al.*, 2012; Bulgarelli *et al.*, 2013; Horton *et al.*, 2014; Coleman-Derr *et al.*, 2016; Wagner *et al.*, 2016), bacterial communities were largely dominated by Proteobacteria (> 80%). At the order level, Burkholderiales (29.3%) and Sphingomonodales (27.9%) were dominant (Figure 2a). In comparison with autumn, samples collected in spring were enriched for Burkholderiales and depleted for Sphingomonodales in the root compartment (Figure 2a). Germination timing in autumn did not impact the relative abundance of bacterial order in plants collected in spring (Figure 2a).

**Figure 2.**
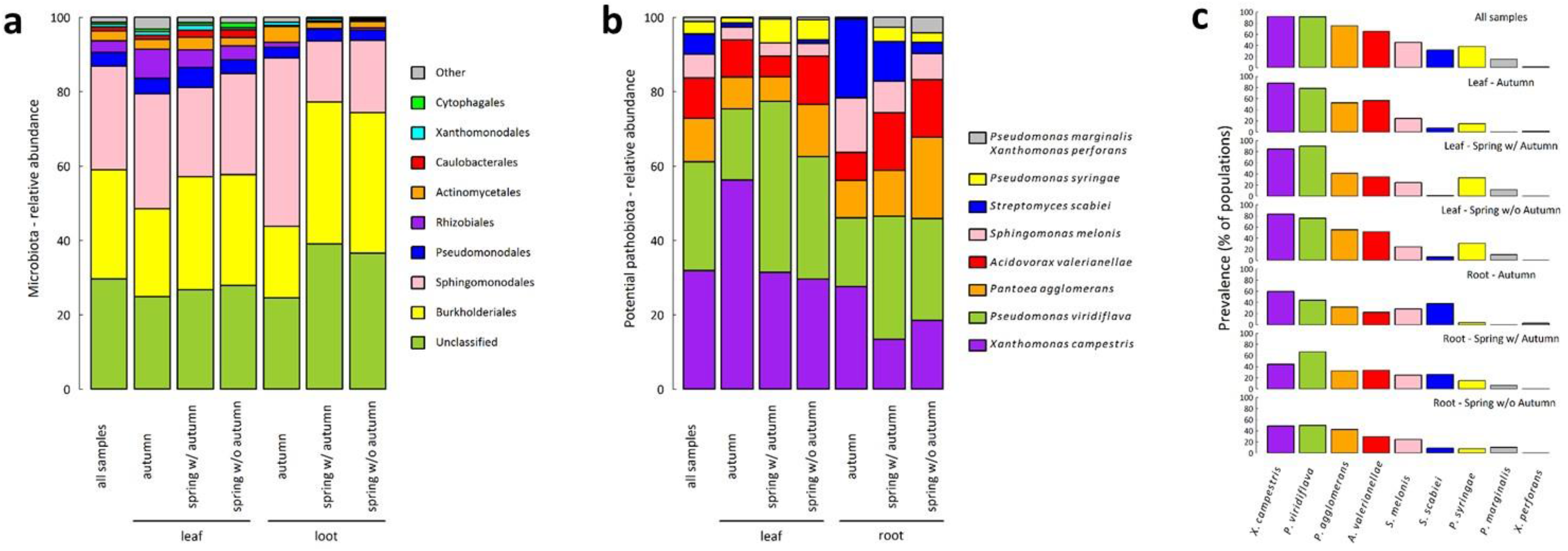
Stacked bar plots of the relative abundances of the major bacterial phyla and bacterial species for both microbiota and potential pathobiota of the *A. thaliana* populations collected in the Midi-Pyrénées region. (**a**) Stacked barplots representing the relative abundance for the ten most abundant bacterial orders of the microbiota. ‘All samples’ n =1655; Leaf: ‘autumn’ n= 314, ‘spring w/ autumn’ n = 245, ‘spring wo/ autumn’ n = 262; Root: ‘autumn’ n= 309, ‘spring w/ autumn’ n = 267, ‘spring wo/ autumn’ n = 258. In comparison with autumn, samples in spring were enriched for Burkholderiales (χ^2^ = 7.93, *P* < 0.01) and depleted for Sphingomonodales (χ^2^ = 18.18, *P* < 0.001) but only in the root compartment. (**b**) Stacked barplots representing the relative abundance of the nine bacterial species representing the potential pathobiota. ‘All samples’ n = 1203; Leaf: ‘autumn’ n = 273, ‘spring w/ autumn ‘ n = 209, ‘spring wo/ autumn’ n = 213; Root: ‘autumn’ n= 180, spring w/ autumn’ n = 171, ‘spring wo/autumn’ n = 157. For the seasonal groups ‘autumn’ and ‘spring w/autumn’, we observed a depletion of *X. campestris*(‘autumn’, χ^2^ = 15.67, *P*<0.01; ‘spring with autumn’, χ^2^ = 8.26, *P*< 0.05) and an enrichment of *S. scabiei*(‘autumn’, χ^2^ = 18.61 *P*<0.001; ‘spring with autumn’, χ^2^ = 9.31,*P* < 0.05) in the root compartment in comparison with the leaf compartment. In the leaf compartment, an enrichment of *P. viridiflava* between autumn and spring (χ^2^ =15.20, *P*< 0.01) was associated with a depletion of *X. campestris*(χ^2^= 11.50,*P*<0.01). A correction for the number of tests was performed to control the False Discovery Rate at a nominal level of 5%. c) Prevalence (number of populations in which the potential pathogen species was detected) of the nine bacterial species characterizing the potential pathobiota.

Across all samples, we identified 9 bacterial species described as phytopathogenic by the International Society of Plant Pathology Committee on the Taxonomy of Plant Pathogenic Bacteria (Bull *et al.*, 2010, 2014) (Figure 2b). Among them, the three most abundant species representing more than 72% of the whole potential pathobiota were *X. campestris* (31.9%), *P. viridiflava*(29.2%) and *P. agglomerans* (11.7%) (Figure 2b). For the seasonal groups ‘autumn’ and ‘spring w/autumn’, we observed a depletion of *X. campestris* and an enrichment of *S. scabiei* in the root compartment in comparison with the leaf compartment (Figure 2b). In the leaf compartment, an enrichment of *P. viridiflava* between autumn and spring was associated with a depletion of *X. campestris* (Figure 2b). Similarly to the microbiota, no effect of germination timing in autumn was observed on the relative abundance of the potential pathogenic species for plants collected in spring (Figure 2b).

It is noteworthy that the relationship between the total relative abundance among samples and the prevalence among populations was weak for some potential pathogenic bacterial species (Figure 2c). For example, although the total relative abundance of *P. syringae* and *S. scabiei* among all samples ranged from 3.4% to 5.5% respectively (Figure 2b), these two species were present on average in more than 31% of the populations (Figure 2c). Visual inspection of original data suggests that this pattern is mainly explained by the presence of few highly infected plants in many populations.

To confirm the pathogenic behaviour of the potential pathobiota, representative bacterial strains of the three most abundant species were tested for their pathogenicity on their natural host *A. thaliana* and on a non-host plant (tobacco). For the *P. syringae* complex (including both *P. syringae sensu stricto* and *P. viridiflava*; Supplementary Text), we isolated 97 strains (74 *P. viridiflava* strains and 23 *P. syringae sensu stricto* strains; Supplementary Text, Supplementary Data 3, Supplementary Figure S4). Their pathogenicity was assessed based on *in planta* bacterial growth and disease symptoms in *A. thaliana* and a Hypersensitive Response (HR) test on tobacco (Supplementary Text). We found that: i) 84 strains of the *P. syringae* complex induced a HR on tobacco (Supplementary Text, Supplementary Data 3), ii) all the four strains tested for *in planta* growth were able to reach a population size of 10^6^ CFU.cm^−2^ 7 days post inoculation on *A. thaliana* (Supplementary Figure S5a, Supplementary Table S2), and iii) seven out of eight strains tested were able to induce disease on at least one of the eight *A. thaliana* local accessions tested (Supplementary Text, Supplementary Figure S5b, S6, S7, Supplementary Table S3). For *X. campestris*, 62 strains were isolated (Supplementary Text, Supplementary Figure S8) and all of them induced disease symptoms on the *A. thaliana* Kas-1 accession (Supplementary Figure S9), which is susceptible to most *X. campestris* strains isolated from crops (Huard-Chauveau *et al.*, 2013). In addition, 58 of the 62 *X. campestris* induced a HR on tobacco (Supplementary Text, Supplementary Data 3). For *P. agglomerans*, we isolated a single strain (Supplementary Text, Supplementary Figure S10) that was able to induce disease symptoms on all the 23 *A. thaliana* local accessions tested (Supplementary Text, Supplementary Figure S11, S12, Supplementary Table S4). Taken together, these results support that most of the strains identified here as part of the *A. thaliana* potential pathobiota have a pathogenic behavior on *A. thaliana*. In addition, the relative abundance of the potential pathobiota was significantly higher in *A. thaliana* individuals with visible disease symptoms (4.5%) than in asymptomatic *A. thaliana* individuals (1.6%) when sampled *in situ* (general linear model, *F* = 26.05, *P* < 0.001) (Supplementary Figure S13), strengthening the potential pathogenic behavior of the pathobiota identified in this study.

### Alpha-diversity of the *A. thaliana* microbiota and potential pathobiota

We investigated whether the level of α-diversity (species richness and Shannon index) estimated for each sample was driven by the effects of season, plant compartment and population. Statistical results globally led to similar biological conclusions between species richness and Shannon index (Supplementary Tables S5 - S12). Bacterial communities of the microbiota were on average less diverse in roots than in leaves, in particular in autumn (Figure 3a, Supplementary Tables S5, S6, S7). For the potential pathobiota, a similar pattern was observed in autumn but not in spring where the potential pathogenic communities were as diverse in roots as in leaves (Figure 3b, Supplementary Tables S5, S6, S7). More importantly, strong differences among populations in the dynamics of α-diversity between autumn and spring were observed for the microbiota and to a lesser extent for the potential pathobiota (Figure 4, Supplementary Tables S5, S6).

**Figure 3.**
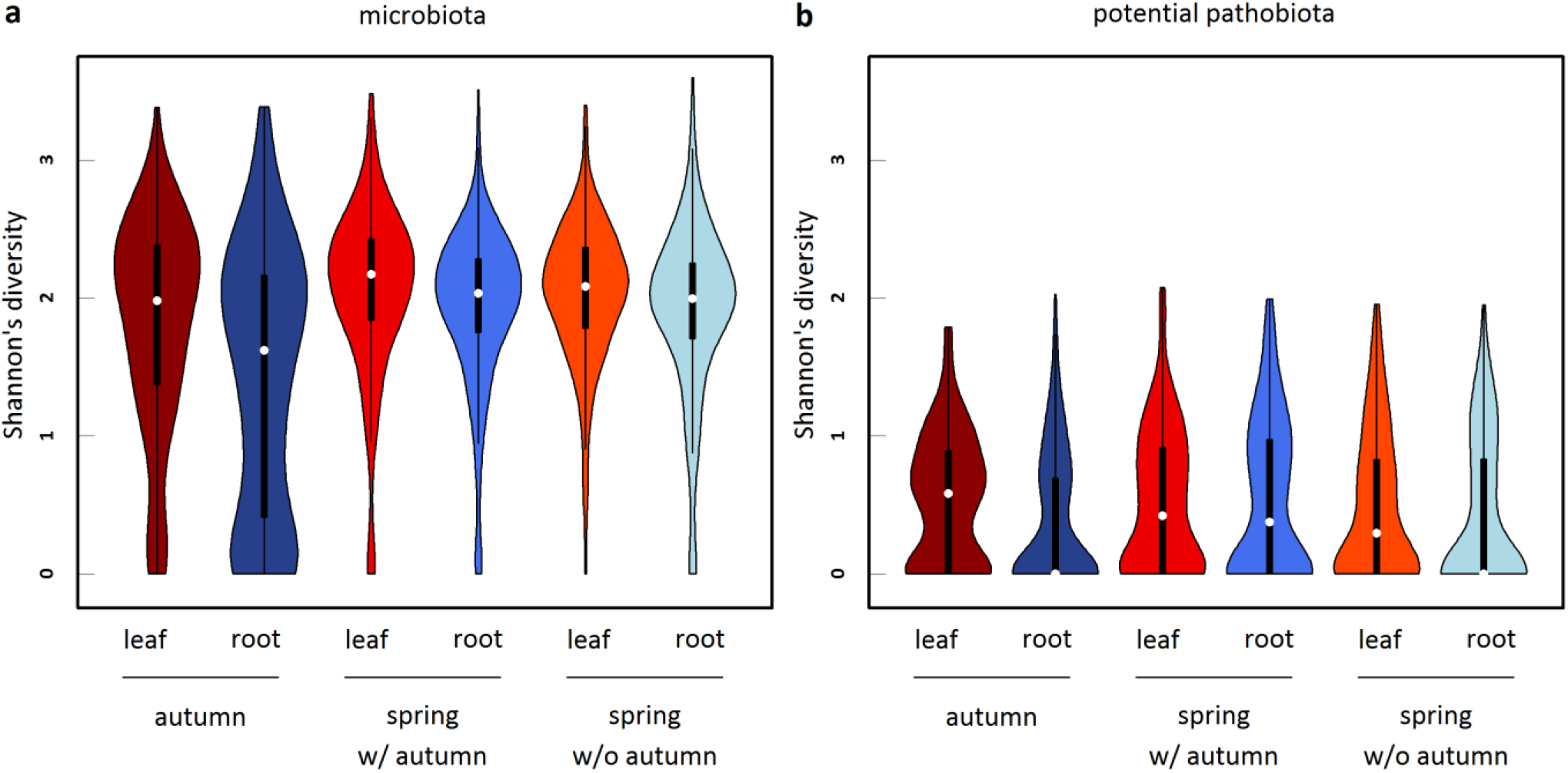
Violin plots (i.e. box-and-whisker plot overlaid with a kernel density plot) representing the seasonal variation of the α-diversity inferred as Shannon’s index for both (**a**) microbiota and (**b**) potential pathobiota. Leaf and root samples are represented with a red and blue color scale, respectively. Microbiota: ‘Autumn – Leaf’ n= 314, ‘Autumn – Root’ n = 309, ‘Spring w/ Autumn - Leaf’ n= 245, ‘Spring w/ Autumn - Root’ n= 267,‘Spring w/o Autumn - Leaf’ n= 262, ‘Spring w/o Autumn - Root’ n= 258. Pathobiota: ‘Autumn – Leaf’ n= 273, ‘Autumn – Root’ n = 180, ‘Spring w/ Autumn - Leaf’ n= 209, ‘Spring w/ Autumn - Root’ n= 171,‘Spring w/o Autumn - Leaf’ n= 213, ‘Spring w/o Autumn - Root’ n= 157.

**Figure 4.**
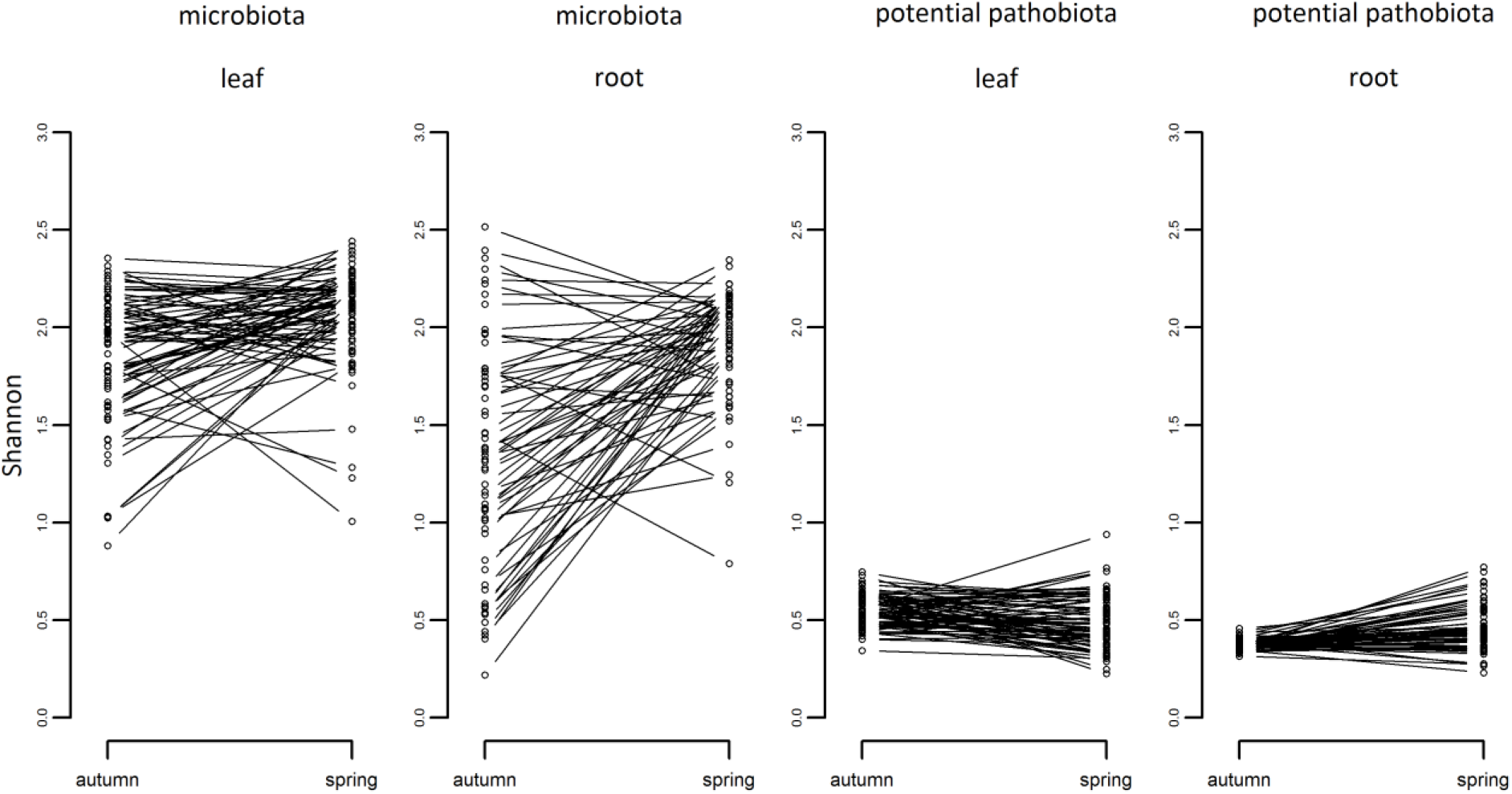
Variation among populations in the dynamics of α-diversity between autumn and spring. Each dot corresponds to the mean α-diversity (estimated as BLUPs) of a population. ‘leaf’ n = 74 populations, ‘root’ n = 62 populations.

Whatever the seasonal group considered, variation in α-diversity of the microbiota was first explained by differences among populations (∼16.8% for species richness and ∼26.2% for Shannon index) (Figure 4, Supplementary Tables S8-S11). In contrast, the main source of variation in the α-diversity of the potential pathobiota largely differs between the two seasons (Supplementary Tables S8-S11). Variation in α-diversity of the pathobiota in autumn and spring was first explained by the factor ‘plant compartment’ (∼12.5%) and the factor ‘population’ (∼13.8%), respectively (Supplementary Table S11). At the ‘plant compartment × seasonal group’ level, the level of differentiation among populations for Shannon index was on average almost twice higher for the microbiota than for the potential pathobiota (∼38.3% *vs*∼14.5% of variance explained by the factor ‘population’) (Supplementary Table S12).

No effect of germination timing in autumn was observed on the microbiota and pathobiota α-diversity of plants collected in spring (Supplementary Table S7, S11).

### Relationships between microbiota and potential pathobiota α-diversity: testing for the invasion paradox

By considering all samples, we observed a highly significant humped-back relationship between the species richness of the potential pathobiota and the species richness of the microbiota (Figure 5a, Supplementary Table S13, Supplementary Figure S14). This humped-back relationship was robust whatever the ‘seasonal group × plant compartment’ considered (Figure 5b, 5c, Supplementary Table S13). A similar humped-back relationship was observed when considering Shannon index instead of species richness, with the exception of the root compartment in the seasonal group ‘spring w/ autumn’ (Figure 5d, 5e, 5f, Supplementary Table S13). This humped-back relationship indicates that a poorly diversified potential pathobiota was associated with either a highly or a poorly diversified microbiota, whereas a highly diversified potential pathobiota was found in presence of microbiota with an intermediate level of diversity. Altogether, these results are in line with theoretical expectations of the invasion paradox (Supplementary Figure S1).

**Figure 5.**
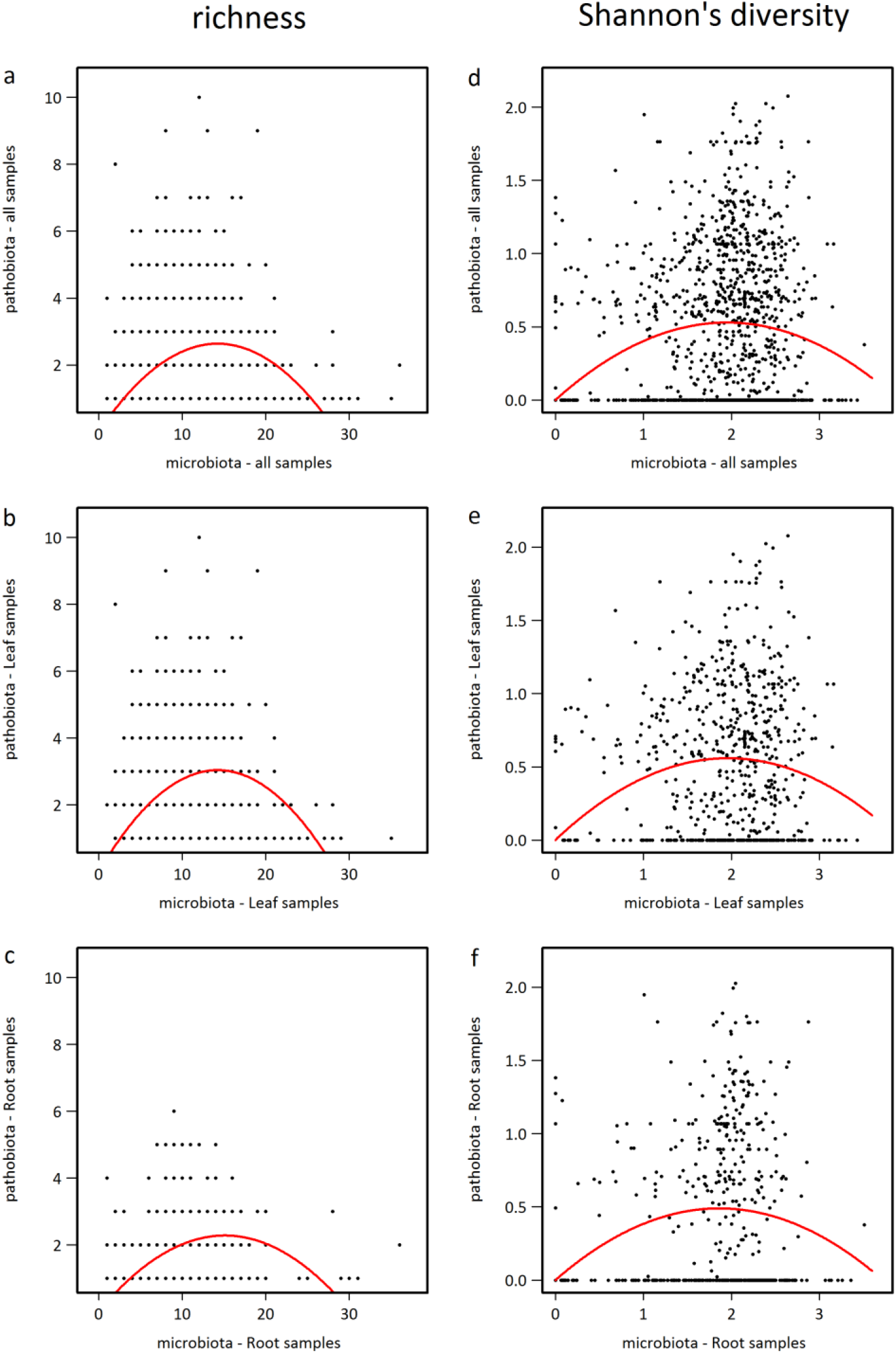
Humped-back relationships between potential pathobiota α-diversity and microbiota α-diversity. (**a**) Species richness when considering all samples. (**b**) Species richness when considering samples from the leaf compartment. (**c**) Species richness when considering samples from the root compartment. (**d**) Shannon’s index when considering all samples. (**e**) Shannon’s index when considering samples from the leaf compartment. (**f**) Shannon’s index when considering samples from the root compartment. The red lines indicate a significant quadratic relationship, according to the following non-linear model: pathobiota’s diversity ∼ k^*^microbiota’s diversity – q^*^microbiota’s diversity^*^microbiota’s diversity.

### Composition and structure of *A. thaliana* microbiota and potential pathobiota

For the microbiota, the first two PCoA axes explained 20% of the β-diversity (Figure 6). The microbiota was structured according to a pattern of two perpendicular branches (Figure 6a). The abundance variation of the most abundant OTU, i.e. an OTU belonging to the genus *Sphingomonas*, explained up to 20.7% of the variation along the first branch (Figure 6b). The abundance variation of the second most abundant OTU (an OTU belonging to the family Oxalobacteraceae), was related to the variation along the second branch (Figures 6c). For the potential pathobiota structured according to a pattern of three branches (Figure 6d), the first two PCoA axes explained 37% of the β-diversity (Figure 6d). The variation along two of these branches was related strongly but independently to the abundance variations of *P. viridiflava* and *X. campestris* (Figures 6e, 6f).

**Figure 6.**
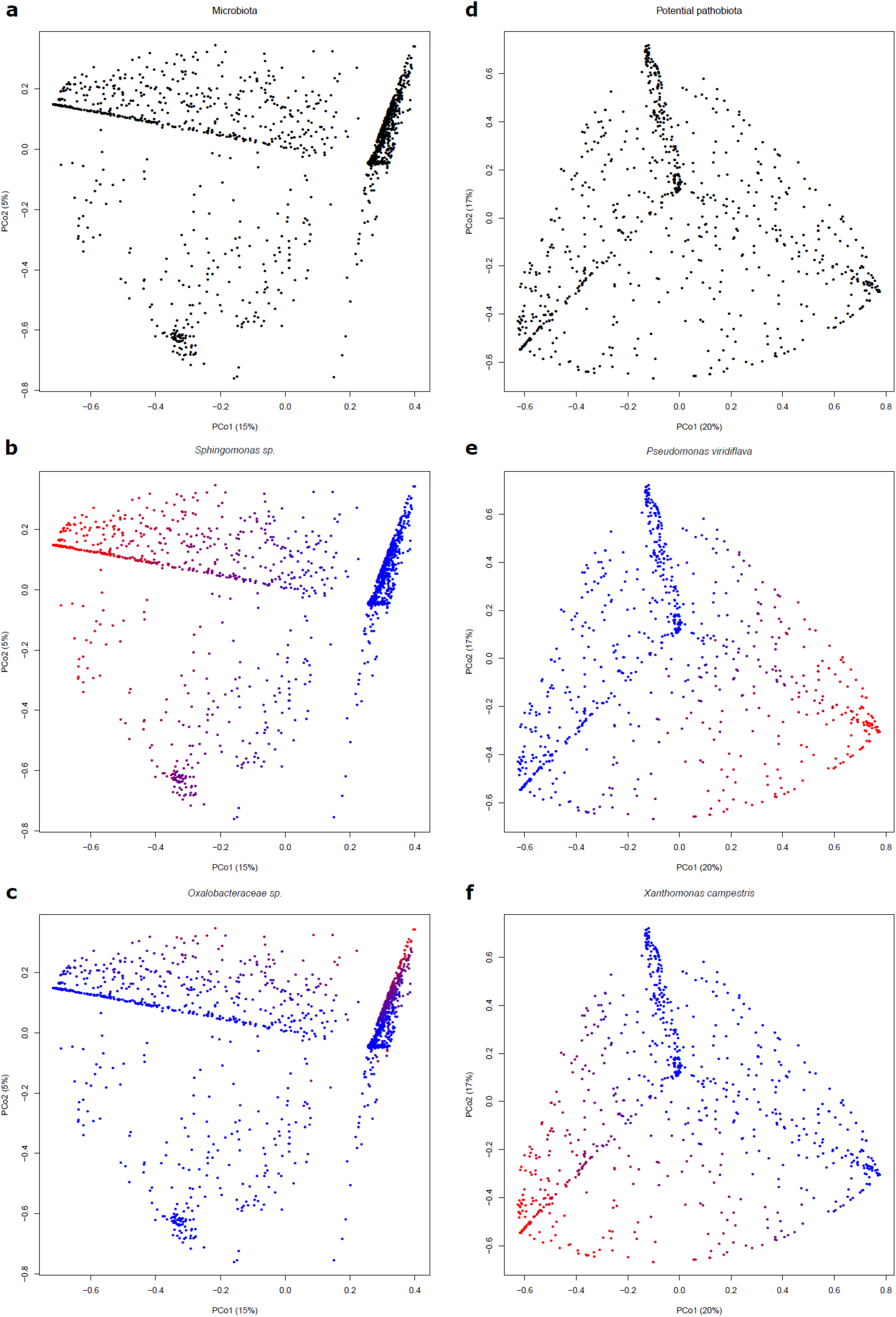
Bacterial composition and structure of *A. thaliana* illustrated by Principal coordinates (PCoA) plots of Hellinger dissimilarity matrices for microbiota (left panels) and potential pathobiota (right panels). n = 1655 samples for the microbiota matrix and n = 1203 for the potential pathobiota matrix. (**a**) PCoA plot of microbiota for all samples indicates that bacterial composition is mainly structured along two axes. The (**b**) and (**c**) plots indicate the relationships between the microbiota composition and the abundance of the two most abundant OTUs, i.e. *Sphingomona ssp.*, and *Oxalobacteraceae unclassified sp.*, respectively. (**d**) PCoA plot of potential pathobiota for all samples indicates that bacterial composition is mainly structured along three axes. The (**e**) and (**f**) plots indicate the relationships between the potential pathobiota composition and the abundance of the two most abundant potential pathogenic OTUs, i.e. *Pseudomonas viridiflava* and *Xanthomonas campestris*, respectively. Red-to-blue color gradient indicates high-to-low OTU abundance.

We observed strong differences among populations for the dynamics of β-diversity of the microbiota between autumn and spring, with the factor ‘season × population’ explaining 53.8% and 43.2% of the variation along the first and second PCoA axes, respectively (Supplementary Figure S15, Supplementary Tables S5,S6). The dynamics of β-diversity of the potential pathobiota between autumn and spring was also dependent on the considered population but to a much lesser extent than what was observed for the microbiota β-diversity (i.e. 9.8% and 7.6% of the variation along the first and second PCoA axes, respectively) (Supplementary Tables S5, S6).

Within each seasonal group, the variation of microbiota along the PCoA axes was largely explained by differences among populations (up to ∼81%) (Supplementary Figure S16, Supplementary Tables S8 – S11), while the variation of the potential pathobiota β-diversity was first explained by the factors ‘population’ (∼16.2%) and ‘plant compartment’ (∼12.6%) when considering the first and second PCoA axis, respectively (Supplementary Table S11). At the ‘plant compartment × seasonal group’ level, the level of differentiation among populations for β-diversity was on average higher for the microbiota than for the potential pathobiota (∼76.5% *vs* ∼10.8% of the first PCoA axis variance explained by the factor ‘population’) (Supplementary Table S12, Supplementary Figure S16).

An effect of germination timing in autumn on the β-diversity of plants collected in spring was not observed on microbiota but on potential pathobiota. In the leaf compartment, variation along the first PCoA axis was driven by differences among populations with early autumn germinants (i.e. ‘spring with autumn’ season group), while variation along the second PCoA axis was driven by differences among populations with late autumn germinants (i.e. ‘spring without autumn’ season group) (Supplementary Table S12, Supplementary Figure S16).

### Relationship between microbiota and potential pathobiota β-diversity

For each ‘season group × plant compartment’ combination, a non-negligible percentage of variation of the potential pathobiota composition (16.9% on average) was explained by a minor fraction of the variation of the microbiota composition (3.3% on average) (Figure 7). The identity of the microbiota OTUs associated with variation of the potential pathobiota composition largely differs between seasons, plant compartment (in particular in autumn) and germination cohorts in autumn (Figures 7 and 8). Among the 17 candidate OTUs of the microbiota, the fifth most abundant microbiota OTU (*Pseudomonas moraviensis*) was the most prevalent microbiota OTU across the six ‘season group × plant compartment’ combinations. However, a large fraction of the candidate microbiota OTUs (58.8%) corresponds to unclassified OTUs at the order level.

**Figure 7.**
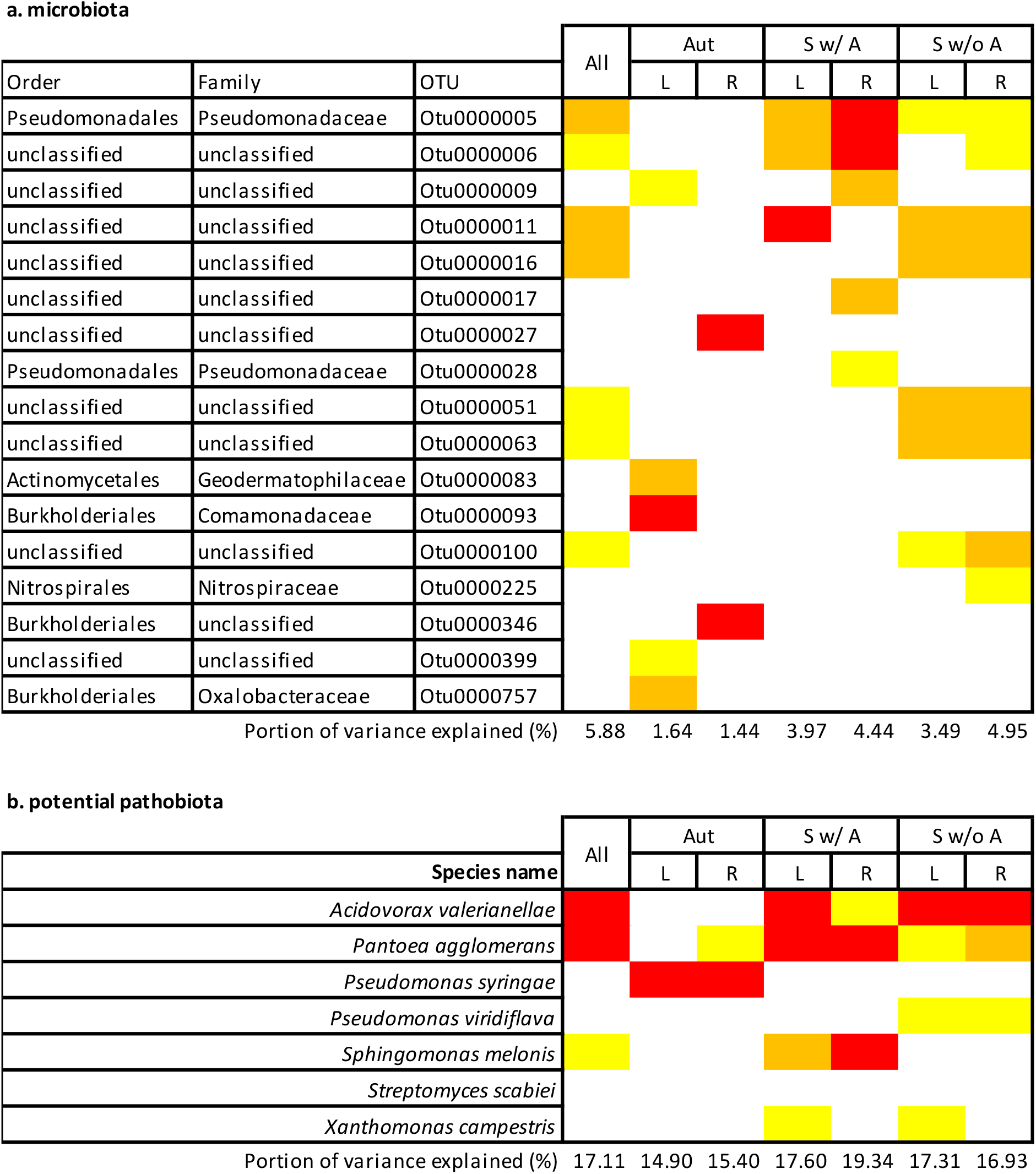
Relationships between microbiota β-diversity and potential pathobiota β-diversity based on a Sparse Partial Least Square Regression (sPLSR). Only OTUs with a loading value above 0.2 in more than 75% of the 1,000 Jackknife resampled matrices were considered as significant. The color gradient indicates the strength of the loading values for both microbiota OTUs and potential pathobiota OTUs (yellow: 0.2 < loadings < 0.35, orange: 0.35 < loadings < 0.5, red: loadings > 0.5). ‘All’, ‘Aut’, ‘Sw/ A’ and ‘Sw/o A’ stand for all samples and samples from the three seasonal groups (i.e. ‘autumn’ populations, ‘spring w/ autumn’ populations and ‘spring w/o autumn’ populations), ‘L’ and ‘R’ stand for leaf and root, respectively. ‘All’ n = 1655; ‘Aut - L’ n= 314, ‘Sw/ A - L’ n = 245, ‘Sw/o A - L’ n = 262; ‘Aut – R’ n= 309, ‘Sw/ A - R’ n = 267, ‘Sw/o A – R’ n = 258.

**Figure 8.**
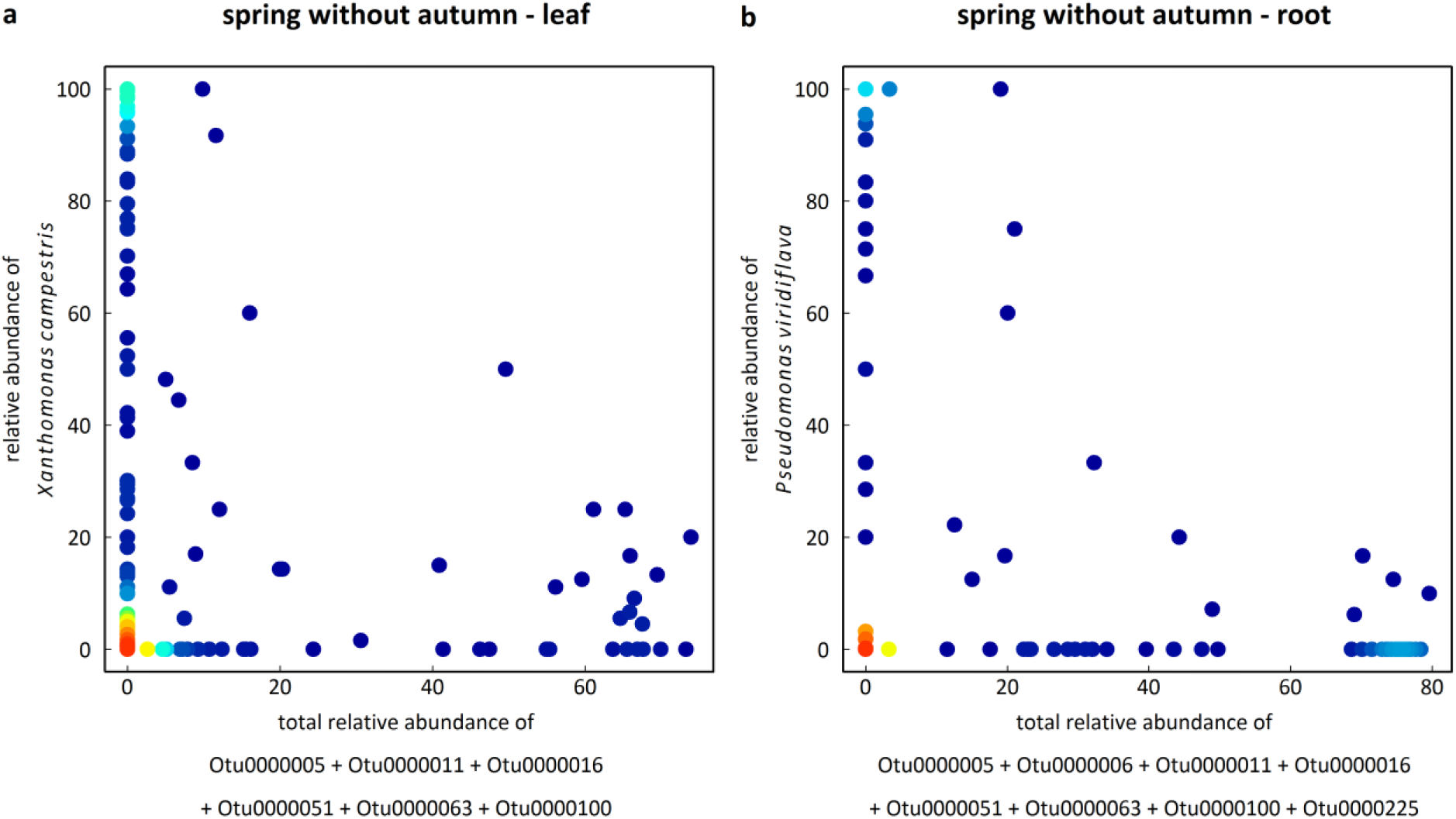
2D density plots illustrating the relationships between the relative abundance of the two most abundant bacterial species of the potential pathobiota and the total relative abundance of the microbiota OTUs identified by sPLSR. (**a**) Example with *X. campestris* in the seasonal group ‘spring with autumn’ in the leaf compartment (Pearson’s *r* = −0.15, *P* = 0.0187). (**b**) Example with *P. viridiflava* in the seasonal group ‘spring with autumn’ in the root compartment (Pearson’s *r* = −0.17, *P* = 0.0006). Red-to-blue color gradient represents high-to-low density gradient.

## DISCUSSION

### The characterization of the *A. thaliana* potential pathobiota in an ecological context

Several evidences suggest that the potential pathobiota was well characterized in our set of 163 natural populations of *A. thaliana*, thereby allowing investigating the bacterial microbiota-potential pathobiota relationships. First, according to the *gyrB* community profiling, *X. campestris* and *P. viridiflava* were the two most abundant species of the *A. thaliana* potential pathobiota. These observations are congruent with a previous bacterial community profiling obtained with a 16S rRNA region of 1.4kb from a field experiment on wild-type and mutant lines of *A. thaliana* (Kniskern *et al.*, 2007). Second, we detected *in situ* a strong relationship between the presence of disease symptoms and the relative abundance of the potential pathobiota. Third, pathogenicity tests on both host and non-host plants confirmed that most of the strains belonging to the three most abundant species - composing almost three quarters of the whole potential pathobiota - have a pathogenic behavior. Finally, we found a significant and strong positive relationship between relative abundance and α-diversity of the pathobiota (Supplementary Table S14). The latter result suggests that plants are more often infected by consortia of pathogenic species than by a single pathogenic lineage, which is in line with similar observations obtained in humans and animals (Faust *et al.*, 2012). For example, multiple closely related *Borrelia* genospecies have been found to display a positive co-occurrence in ticks (Herrmann *et al.*, 2013). Studies on plant pathogens also suggest that co-infection is a frequent process mediated by niche functionality of the colonizing species (Lamichhane and Ventury, 2015). More broadly, primary infecting pathogenic lineages can open the route for opportunistic lineages, as previously demonstrated for brassica black rot disease in which leaf infection by *Xanthomonas* is followed by the infection of a soft-rotting bacterium responsible for the rotting disease (Williams, 1980).

### Putting the characterization of bacterial communities in an ecological genomics framework

In this study, we found that the *in situ* microbiota and potential pathobiota of *A. thaliana* were affected by the combined effects of season, plant compartment and population. In particular, we observed a strong dynamics in community succession of the microbiota between seasons, reinforcing the need to study the dynamics of bacterial communities over the entire plant life cycle (from seed to seed) across a large range of native habitats (Roux and Bergelson, 2016). In addition, our study revealed that seasonal community succession of the microbiota largely differed among the 163 populations. Because α- and β-diversity varied at a very small geographic scale (Supplementary Figure S17), understanding among-population variation of seasonal community succession will require a thorough characterization of abiotic and biotic factors known to influence microbiota variation in plant species, such as soil conditions (Lundberg *et al.*, 2012; Bulgarelli *et al.*, 2013), micro-local climatic conditions (Classen *et al.*, 2015) and plant community composition (Aleklett *et al.*, 2015; Geremia *et al.*, 2016).

In contrast to the microbiota, we found that plant compartment mainly influenced the composition of the potential pathobiota as well as the relative abundance of the most abundant pathogenic species. Our results are in accordance with previous studies reporting that pathogenic species such as *P. syringae sensu lato* and *X. campestris* evolved specific strategies (e.g. entry by stomata, hydatodes and wounds) to infect the leaf compartment in a wide range of crops (Mansfield *et al.*, 2012). On the other hand, the most abundant OTUs of the microbiota were shared between leaves and roots. In agreement, the use of a gnotobiotic *A. thaliana* plant system allows to demonstrate potential reciprocal relocation between root and leaf microbiota members (Bai *et al.*, 2015). In the same study, whole-genome sequencing and functional analysis of bacteria associated with both leaves and roots of *A. thaliana* highlighted a clear taxonomy and functional overlap of the bacterial populations inhabiting this plant species (Bai *et al.*, 2015). Altogether, these results are in contrast with previous studies on human microbiota demonstrating a remarkable organization of microbes into body site niches (Faust *et al.*, 2012; Belkaid and Hand, 2014). This discrepancy might originate from a lack of specialized plant tissues constituting strictly specialized niches for commensal bacteria as observed in human organs.

### The invasion paradox is mediated by distinct microbiota composition between seasons and plant organs

In agreement with theoretical expectations on the invasion paradox (Levine and D’Antonio, 1999; Fridley *et al.*, 2007; Tomasetto *et al.*, 2013), we observed a humped-back relationship between potential pathobiota α-diversity and microbiota α-diversity. Such a pattern may have been observed due to the large range of habitats where *A. thaliana* plants have been collected, thereby increasing the range of dimensionality of *in planta* ecological niches available for microbes. Besides the interplay between diversity and niche dimensionality, other hypotheses can contribute to the explanation of the humped-back relationship. An increase in microbiota diversity can be associated with an increase of antagonistic (e.g. production of anti-microbial compounds) and predator (e.g. grazing) species (Mallon *et al.*, 2015b), leading to a negative relationship between potential pathobiota α-diversity and microbiota α-diversity. Concerning the positive relationship between poor diversified potential pathobiota and microbiota, several non-exclusive hypotheses can be advanced. Firstly, ecological disturbance can increase the random establishment of a single pathogenic species with negative consequences on native species diversity (Kinnunen *et al.*, 2016). Secondly, by the producing virulence proteins, a pathogen can exclusively invade a given plant compartment (Hacquard *et al.*, 2017). Thirdly, both microbiota and potential pathobiota can be poorly diversified because other microbial communities (such as fungal and oomycete communities) exploit most of the resources available in the plant, leading to niche inter-kingdom competition (Agler *et al.*, 2016).

Interestingly, the pattern of invasion paradox was robust between compartments but also between seasons, despite a strong seasonal community succession of the microbiota in most populations. While this observation reinforces the importance of α-diversity in *A. thaliana* populations to maintain a microbiota balance to prevent pathogen occurrence, it also suggests that the potential pathobiota composition was associated with season-specific combinations of microbiota OTUs. Accordingly, the identity of the microbiota OTUs associated with variation of the potential pathobiota composition largely differs between seasons. This dynamics in the potential biomarkers (i.e. specific bacteria taxa preventing pathogen spread) controlling pathogen’s invasion may result from niche overlap and other mechanisms regulating microbe-microbe interactions that drastically influence the bacterial composition. A better understanding of the processes underlying the observed invasion paradox will require the isolation of a large number of strains representative of the microbiota found in the 163 *A. thaliana* populations. This step will especially be relevant for the numerous microbiota OTUs that are unclassified at the order level but associated with variation of the potential pathobiota composition.

However, we should acknowledge that a substantial fraction of the variation of the *A. thaliana* pathobiota was not explained by the microbiota variation. This fraction of unexplained pathobiota variation may result from (i) habitat-specific combinations of bacterial species controlling the potential pathobiota, (ii) the control of bacterial pathogen species by fungal or oomycete microbes (Agler *et al.*, 2016), and (iii) the genetics of *A. thaliana* shaping natural variation in both pathogen’s abundance and prevalence (Roux & Bergelson, 2016). Teasing apart the relative roles of these putative factors will require a thorough complementary ecological and genomic characterization of our natural populations of *A. thaliana*.

## ACKNOWLEDGEMENTS

We are also grateful to the staff of the LIPM greenhouse for their assistance during the growth chamber experiments. This work was funded by the Région Midi-Pyrénées (CLIMARES project), the LABEX TULIP (ANR-10-LABX-41, ANR-11-IDEX-0002-02) and the Métaprogramme MEM (INRA, Metabar programme).

## SUPPLEMENTARY FILES

**Supplementary Text**: supplementary material and methods and supplementary results.

**Supplementary Figure S1**. The invasion paradox. (A) Species richness of both native and alien species is positive correlated with the quality of the environment. Abiotic factors often have similar effects on biodiversity and invader success. (B) The increasing quality of the environment leads to a positive relationship between species richness of alien species and species richness of native species. (C) Elton’s hypothesis describing a negative association between the increasing of the biodiversity and the success of the invaders. The interplay between resource availability and diversity in determining invasion resistance leads to the well-known phenomenon called ‘invasion paradox’. (D), (E) and (F) indicate application of the theories to microbial communities.

**Supplementary Figure S2**. Distribution of the number of reads (expressed in log_10_) for the 6,627 OTUs obtained after data filtering (top panel) and for the remaining 271,706 OTUs discarded with the filtering (bottom panel). The mean number of reads per OTU maintained after filtering was 51.3 fold higher than the one of the discarded OTUs.

**Supplementary Figure S3**. PCoA perfomed on a Hellinger distance matrix based on rarefied data (top panel) and on a Jaccard similarity coefficient matrix distance (bottom panel).

**Supplementary Figure S4**. Phylogenetic tree inferred with a Neighbor Joining (NJ) model (3000 bootstrap repetitions) and based on the *cts* sequences (350bp) of the 97 *P. syringae* strains isolated from the 163 *A. thaliana* populations. Bootstrap values are shown at each node and names of the strain at each branch. All the strains belong to the *P. syringae* complex and are mainly distributed in the phylogroups 7, 9, 1, 2, 11 and 13. Reference *P. syringae* strains representative of the 13 phylogroups are labeled in green. Phylogroup affiliation was based on the previous work from Berge *et al*. (2014).

**Supplementary Figure S5**. Pathogenicity of *Pseudomonas* sp. strains isolated from the 163 *A. thaliana* populations of the region Midi-Pyrénées. (A) Mean bacterial growth across eight accessions collected in the region Midi-Pyrénées for two strains of *P. viridiflava* (0114-Psy-NAUV-BL and 0124-Psy-SAUB-AL) and two strains of *P. syringae sensu stricto* (0132-Psy-BAZI-AL and 0143-Psy-THOM-AL). (B) Illustration of symptoms observed three days after inoculation for two strains of *P. syringae*. Presence and absence of symptoms was observed on each of the eight accessions collected in the region Midi-Pyrénées for the strains 0111-Psy-RAYR-BL and 0117-Psy-NAZA-AL, respectively. D0, D3, D5 and D7 indicate the number of days after inoculation.

**Supplementary Figure S6**. Genetic variation among five accessions from the Midi-Pyrénées region for the response to the *P. viridiflava* strain 0124-Psy-SAUB-AL, three days after infiltration. Mock: infiltration with water.

**Supplementary Figure S7**. Heatmap for the symptoms observed three days after inoculation with *P. syringae* strains. The heatmap shows the interactions between eight natural accessions of *A. thaliana* from the region Midi-Pyrénées and eight natural strains belonging to the *P. syringae* complex collected in the same populations than the eight natural accessions.

**Supplementary Figure S8**. Phylogenetic tree inferred with a Neighbor Joining (NJ) model (3000 bootstrap repetitions) and based on the *gyrB* sequences (280 bp) of the 62 *Xanthomonas campestris* strains isolated from the 163 *A. thaliana* populations. Bootstrap values are shown at each node and names of the strain at each branch. All the strains belong to the *X. campestris* group. Reference *Xanthomonas* strains are labeled in blue. Sequences for the reference strains were downloaded on GenBank.

**Supplementary Figure S9**. Distribution of disease symptoms observed on the accession Kas-1 among the 62 *Xanthomonas campestris* strains isolated from the 163 *A. thaliana* populations. The arrows indicate disease symptoms for the accessions Col-0 and Kas-1 and the mutant *rks1-1* with the *X. campestris* control strain *Xcc*568 (Huard-Chauveau *et al.*, 2013).

**Supplementary Figure S10**. Phylogenetic tree inferred with a Neighbor Joining (NJ) model (3000 bootstrap repetitions) and based on the *gyrB* sequences (290 bp) of the *Pantoea agglomerans* strain (labeled in green) isolated from the MONTI-DL population. Bootstrap values are shown at each node and names of the strain at each branch. All the strains belong to the *X. campestris* group. Reference *Pantoea* sequences were obtained from JGI.

**Supplementary Figure S11**. Pathogenicity of a strain of *P. agglomerans* (0001-Pag-MONTI-DL) isolated from the MONTI-D population of *A. thaliana* population located in the Midi-Pyrénées region. Left panel: Illustration of symptoms observed five days after inoculation on the reference accession Col-0. The two leaves on the left were inoculated on both sides of the midrib while only one side of the midrib has been inoculated for the three leaves on the right. Right panel: genetic variation observed among natural accessions collected in the Midi-Pyrénées region. Symptoms showed in the picture were obtained five days after inoculation of the *P. agglomerans* strain on three *A. thaliana* accessions of the Midi-Pyrénées region (MONTI-A-17, NAUV-B-14 and VILLEM-A-12). As illustrated, the 0001-Pag-MONTI-DL was able to induce disease symptoms but with different severity among the three accessions.

**Supplementary Figure S12**. Dynamics of disease symptoms across 23 accessions of the region Midi-Pyrénées inoculated with the strain 0001-Pag-MONTI-DL of *P. agglomerans*. dpi: days post-inoculation.

**Supplementary Figure S13**. Box-and-whisker plot illustrating the significant difference of the relative abundance of the pathobiota in the leaf compartment between plants with visible disease symptoms - represented by the blue box (n = 424 samples) - and asymptomatic plants-represented by the red box (n = 396 samples) - when sampled *in situ*.

**Supplementary Figure S14**. Comparison of the parameters ‘k’ and ‘q’ of the non-linear model (pathobiota’s diversity ∼ k*microbiota’s diversity – q*microbiota’s diversity*microbiota’s diversity) run on the raw data (red arrows) to a null distribution of those parameters obtained by creating 100 random microbiota OTU matrices paired with 100 random pathobiota OTU matrices.

**Supplementary Figure S15**. Variation among populations in the dynamics of β-diversity (PCoA, first axis) between autumn and spring. Each dot corresponds to the mean β-diversity (estimated as BLUPs) of a population. ‘leaf’ n = 74 populations, ‘root’ n = 78 populations.

**Supplementary Figure S16**. Percentage of variance of the β-diversity among populations. Red and blue bars correspond to microbiota and potential pathobiota, respectively. A correction for the number of tests was performed to control the FDR at a nominal level of 5%: ^ns^ non-significant, ^**^ 0.01>*P*>0.001, ^***^ *P* < 0.001.

**Supplementary Figure S17**. Geographic variation of α-diversity (inferred as Shannon’s index) and β-diversity (when considering the first axis of the PCoA) for the microbiota. For each ‘seasonal group x plant compartment’ combination, blue, green, yellow and red dots correspond to populations from the 1^st^, 2^nd^, 3^rd^ and 4^th^ quartiles of either the α-diversity of the β-diversity distribution (based on population BLUPs). Values indicate the percentage of α-diversity and β-diversity variation among populations. Significance after a FDR correction at a nominal level of 5%: ** 0.01 > *P* > 0.001, *** *P* < 0.001. Number of populations: ‘Autumn – Leaf’ n= 82, ‘Autumn – Root’ n = 82, ‘Spring w/ Autumn - Leaf’ n= 76, ‘Spring w/ Autumn - Root’ n= 79,‘Spring w/o Autumn - Leaf’ n= 80, ‘Spring w/o Autumn - Root’ n= 80.

**Supplementary Table S1**. Names and GPS coordinates (expressed in degrees) of the 163 populations.

**Supplementary Table S2**. *In planta* bacterial growth of four natural *Pseudomonas* strains in four corresponding local natural populations of *A. thaliana*, each represented by two accessions.

**Supplementary Table S3**. Natural interactions between eight local natural *A. thaliana* accessions and eight corresponding local natural *Pseudomonas syringae* strains for disease symptom evolution.

**Supplementary Table S4**. Genetic variation among 23 accessions of the region Midi-Pyrénées for the response to the strain 0001-Pag-MONTI-DL of *P. agglomerans* at four time points after inoculation (dpi, days post-inoculation). *H*^2^: broad-sense heritability.

**Supplementary Table S5**. Natural variation of α-diversity and β-diversity estimators of microbiota and potential pathobiota in natural populations of *A. thaliana* collected in both autumn and spring.

**Supplementary Table S6**. Percentage of variance of α-diversity and β-diversity estimators of microbiota and potential pathobiota explained by the factors ‘seasons’, ‘plant compartment’, ‘population’ and their interactions in natural populations of *A. thaliana* collected both in autumn and spring.

**Supplementary Table S7**. Natural variation of α-diversity and β-diversity estimators of microbiota and potential pathobiota in all the natural populations of *A. thaliana* collected in spring.

**Supplementary Table S8**. Natural variation of α-diversity and β-diversity estimators of microbiota and potential pathobiota in the natural populations of *A. thaliana* collected in the seasonal group ‘autumn’.

**Supplementary Table S9**. Natural variation of α-diversity and β-diversity estimators of microbiota and potential pathobiota in the natural populations of *A. thaliana* collected in the seasonal group ‘spring with autumn’.

**Supplementary Table S10**. Natural variation of α-diversity and β-diversity estimators of microbiota and potential pathobiota in the natural populations of *A. thaliana* collected in the seasonal group ‘spring without autumn’

**Supplementary Table S11**. Percentage of variance of α-diversity and β-diversity estimators of microbiota and pathobiota explained by the factors ‘population’, ‘plant compartment’ and their interactions in natural populations of *A. thaliana* collected either in autumn or ins spring.

**Supplementary Table S12**. Natural variation of α-diversity and β-diversity estimators of microbiota and potential pathobiota for each ‘seasonal group x plant compartment’ combination.

**Supplementary Table S13**. Comparison of the fit between a linear model and a non-linear model on the relationship between the α-diversity estimators (species richness and Shannon index) of the microbiota and the α-diversity estimators (species richness and Shannon index) of the potential pathobiota.

**Supplementary Table S14**. Relationships between the relative abundance of pathobiota and α-diversity estimators (species richness and Shannon index) of the microbiota and potential pathobiota.

**Supplementary Data 1**. Validation of the *gyrB* gene marker.

**Supplementary Data 2**. List of pathogenic bacteria used to determine the potential pathobiota of the 163 *A. thaliana* populations.

**Supplementary Data 3**. Strains belonging to the *P. syringae* complex, *X. campestris* and *P. agglomerans* isolated from the 163 *A. thaliana* populations for the validation of the potential pathobiota. Housekeeping gene sequences used for phylogeny are available in the data set.

**Supplementary Data 4**. Abundance matrix obtained after data filtering for the whole microbiota.

**Supplementary Data 5**. Abundance matrix for the potential pathobiota.

**Supplementary Data 6**. Raw values for both α and β diversity used in the statistical analysis of natural variation of microbiota and potential pathobiota.

